# Insight into perfluorooctanoic acid-induced impaired mouse embryo implantation via single cell RNA-seq

**DOI:** 10.1101/2023.07.02.547373

**Authors:** Xiaoqiang Sheng

## Abstract

PFOA (perfluorooctanoic acid) is a difficult-to-degrade chemical that poses significant risks to human health and the environment. Studies have shown that PFOA affect female reproduction, but effect and mechanism of low doses PFOA expose on endometrial receptivity are unclear. In this study, we found that exposure to low doses of PFOA damaged endometrial receptivity in mice, resulting in decreased embryo implantation rates in mice. Furthermore, using single-cell sequencing technology, we systematically analyzed the specific mechanisms by which PFOA damages endometrial epithelial cell function and the ANGTL signaling pathway between endometrial stromal cells and epithelial cells, leading to embryo implantation failure. The elucidation of this mechanism provides new targets for the treatment of infertility about exposed to PFOA.

## Introduction

Perfluorooctanoic acid (PFOA) is an environmental pollutant that has become widespread worldwide^1^. This chemical is found in a variety of consumer products, such as non-stick cookware, food packaging, and stain-resistant fabrics^2^. PFOA can persist in the environment for years and can bioaccumulate in the food chain^3^. It has been detected in various environmental matrices, such as water, air, soil, and sediment, as well as in the tissues of wildlife and humans. As a result, the general population is frequently exposed to PFOA, and the adverse effects of this exposure on human health are of great concern.

PFOA has been shown to have toxic effects on the human body, including liver damage, immune system dysfunction, developmental and reproductive problems, and an increased risk of certain types of cancer^4^. PFOA exposure is associated with adverse effects on fetal growth and development and has been detected in the breast milk of nursing mothers^5^. The mechanisms by which PFOA causes these health effects are not yet fully understood, but it is believed to disrupt hormone function, damage DNA, and cause oxidative stress.

Studies have shown that PFOA exposure can have negative effects on human reproductive health. It has been reported in the literature that PFOA exposure can cause endocrine disorders^6^. In the female reproductive system, PFOA can damage the mitochondrial electron transport chain of granulosa cells cause abnormal development of ovarian follicles^7^, and cause abnormal gap links between granulosa cells and oocytes^8^. Moreover, abnormal early embryonic development caused by PFOA can be passed on to the next generation^5^. It has been mentioned in the literature that PFOA can reduce the planting rate of adherent spheres in vitro^9^. However, the impact and mechanism of PFOA on endometrial receptivity, a crucial process in embryo implantation, is still unclear.

The endometrium is the inner lining of the uterus, and it undergoes a complex series of changes during the menstrual cycle to prepare for embryo implantation^10^. The implantation of embryos during the early stages of pregnancy involves interactions between the embryo and the endometrial epithelial cells, which line the surface of the endometrium^11^. Recent studies have suggested that endometrial epithelial cells play a crucial role in embryo implantation^12^, and their function can be influenced by exposure to environmental pollutants.

Our experiments found that long-term low-dose PFOA intake will not affect the normal ovulation and pre-implantation embryonic development of the ovary. However, the implantation rate of the embryos was impaired. To understand the underlying mechanisms by which PFOA affects endometrial receptivity, we have employed single-cell sequencing technology to study individual cells and their functions in greater detail. Using single-cell sequencing technology, we have been able to systematically analyze the specific mechanisms by which PFOA damages endometrial epithelial cell function and the ANGTL signaling pathway of endometrial stromal cells and epithelial cells, leading to embryo implantation failure. This information is essential for identifying new therapeutic targets to improve the success rates of in vitro fertilization and embryo transfer (IVF-ET) in PFOA-exposed patients.

## Results

### PFOA exposure impaired mouse embryo implantation

To investigate the effect that PFOA on mouse embryo implantation, we set up a model of PFOA expose with oral ingest 1mg/kg/d (Figure 1A). As shown in Figure 1B-C, there are no significant difference between the Control and PFOA group (27.8±3.8 vs 28.7±3.5, n=6). PFOA expose did not impair the 2-cell, 4-cell, morula, and blastocyst embryo rate (Figure 1D-H). However, the number of implantation embryos (15. 6±1.0 vs 7.8±1.4, n=9) decreased significantly compared to the Control group (Figure I-J).

**Figure 1.**
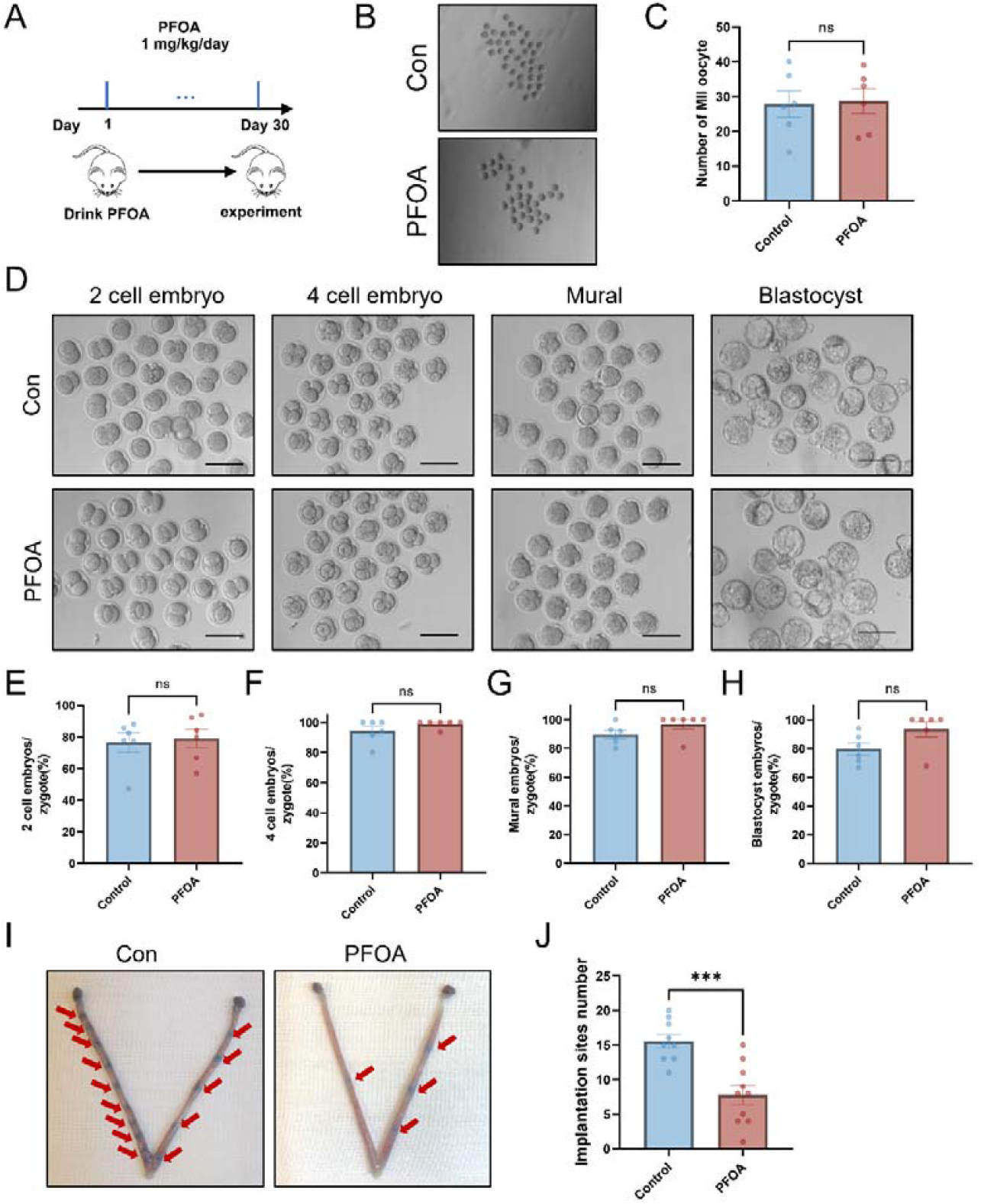
PFOA exposure impaired mouse embryo implantation. (**A**) Strategies for constructing a model of PFOA exposure in mice via drinking water. 1 mg/kg PFOA was added in drinking water of 6-week-old CD-1 mice for 30 days for modeling. (**B-C**) Photographs and numbers of superovulated MII oocytes in Control and PFOA-exposed mice (n=6). (**D**) Photographs showing the stages (2 cell, 4 cell, Mural, and Blastocyst) of embryos from Control and PFOA-exposed mice during IVF. Scale bar = 100 μm. (**E-H**) The two-cell, four-cell, morula and blastocyst rates were analyzed (n=6). (**I**) Uteri from Control and PFOA-exposed mice dpc 4.5 were acquired and stained with Chicago blue dye. Arrow, implantation site. Representative images are shown. (**J**) The number of implantation sites was counted. (***P<0.001, Control (n=9), PFOA (n=10), statistically significant by Student’s t test)

### Single cell transcriptome profiling of PFOA-exposed mice uterus

In order to underline the mechanism of PFOA affect endometrium receptivity, we conducted single cell RNA sequence, bulk-RNA seq, and metabolomic sequencing in dpc 3.5 uterus (Figure 2A). We retained single-cell transcriptomes for 8,093 cells after initial quality controls for uterus from Control and PFOA group (Figure 2B). We identify twelve clusters that were assigned cell identities based on their expression of markers (Figure 2C and Table S1). To further identify the clusters, we assigned the clusters based on known cell-type marker genes. The cell-specific marker genes were specific to stromal cells (*Lum*), epithelial cells (*Krt8*), endothelial cells (*Pecam1*), smooth muscle cells (*Acta2*), immune cells (*Ptprc*), and perivascular (PV) cells (*Rgs5*) (Figure 2D-E). Thus, the clusters were grouped into six main cellular categories: (i) stromal cells, (ii) epithelial cells (iii) endothelial cells, (iv) smooth muscle cells, (v) immune cells, and (vi) PV cells (Figure 2F). To further obtain the dynamic pattern of the gene signature, the six main cluster-specific marker genes were identified (Figure 2G). Among these cell types, the population of epithelial cells decreased markedly when expose to PFOA, which suggest that epithelial cells dysfunction was the main reseason that PFOA exposure impaired mouse embryo implantation (Figure 2H). Our findings provide a resource on various cell types for researchers in this field.

**Figure 2.**
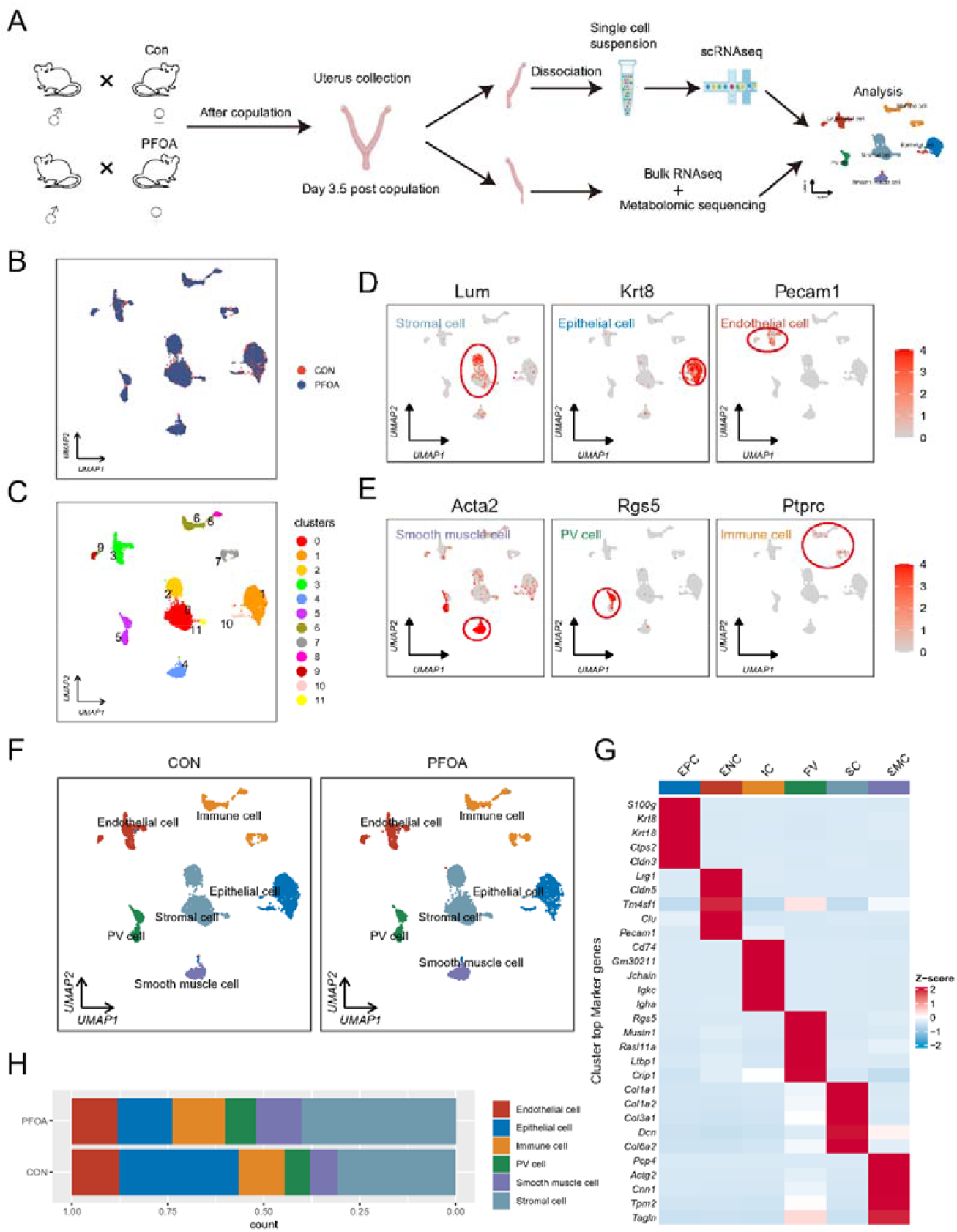
Single cell transcriptome profiling of PFOA-exposed mice uterus. (**A**) Illustration of experimental workflow. Uterus at dpc 3.5 were harvest. Half for single cell RNA sequencing and half for bulk-RNA sequencing and metabolomic sequencing. (**B**) Uniform Manifold Approximation and Projection (UMAP) plot of uterine cell colored by the group (Control and PFOA-exposed). (**C**) UMAP plot of 12 unbiased clusters of combined Control and PFOA-exposed datasets. (**D-E**) Expression profiles of select marker genes in uterine cell projected onto UMAP plots. Lum for Stromal cell, Krt8 for Epithelial cell, Pecam1 for Endothelial cell, Acta2 for Smooth muscle cell, Rgs5 for PV cell, and Ptprc for Immune cell. (**F**) UMAP plots of cells from Control (left) and cells from PFOA-exposed testes (right) colored by the six celltypes. (**G**) Heatmap of top 5 marker genes of the six cell populations. Top 10 marker genes in each cell population are shown in Table S1. EPC: Epithelial cell. ENC: Endothelial cell. IC: Immune cell. PV: PV cell. SC: Stromal cell. SMC: Smooth muscle cell. (**H**) Percentages of the six uterine cell types at Control and PFOA-exposed mice.

### PFOA exposure affect endometrium epithelial cell development

We analysis the bulk RNA seq data in dpc 3.5 uterus (Figure 3A). We identified 596 decreased DEGs and 811 increased DEGs between Control and PFOA group (Figure 3B). Go analysis of down regulated DEGs showed that epithelial cell development and morphogenesis were damaged (Figure 3C). The genes about epithelial cell development (*Myd88, Cldn3, Flnb, Rasn, Notch4, and Ihh)* were showed (Figure 3D). Further qPCR analysis confirmed the results about epithelial cell development genes for *Myd88, Cldn3, Flnb*, and *Rasn* (Figure 3F-I).

**Figure 3.**
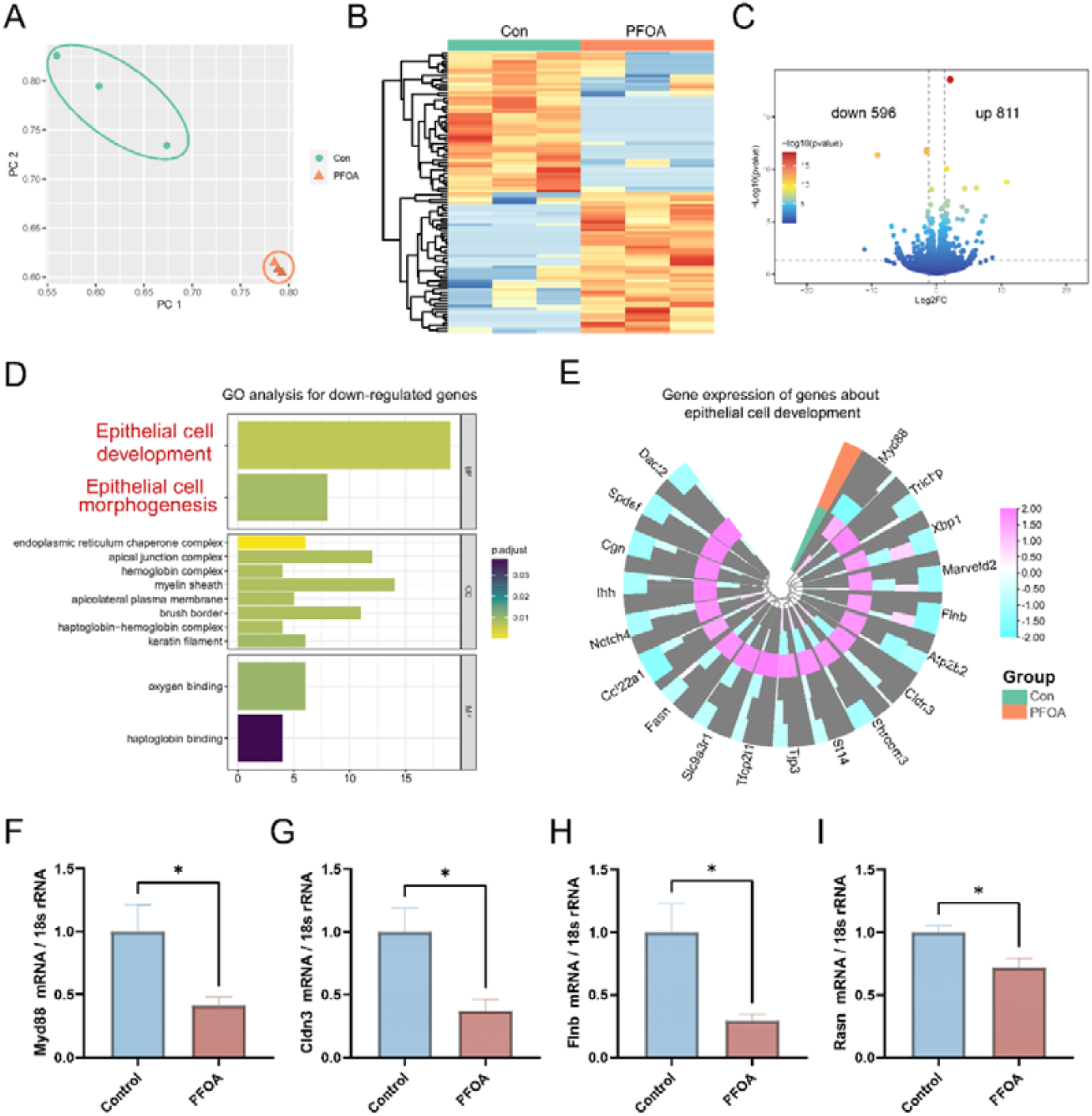
PFOA exposure affect endometrium epithelial cell development. (**A**) Principal component analysis (PCA) analysis of mice uterus in control and PFOA-exposed group. (**B**) Heatmap of differentially expressed genes (DEGs) in control and PFOA-exposed group. (**C**) Volcano map plot of down-regulated and up-regulated genes in control and PFOA-exposed group. Down-regulated genes are 596 and up-regulated genes are 811. (**D**) KEGG pathway analysis of down-regulated genes in the PFOA exposed group. (**E**) Representative genes related to the epithelial cell development pathway expressed in control and PFOA-exposed group. (**F-I**) Real-time PCR analysis of the *Myd88, Cldn3, Flnb*, and *Rasn* transcriptional expression levels in mice uterus (*p < 0.05, statistically significant by Student’s t test).

Furthermore, we analyzed the genetic dynamics of endometrium epithelial cells. When the transcriptome of the epithelial cells were plotted, 5-cell clusters were identified (Figure 4A). As shown in Figure 4B, the cell percent ratio of cluster 2 decreased when expose with PFOA. To examine the features of epithelial cells more closely, we detected the differentially expressed genes in each cluster, and the highly variable expressed genes in epithelial cells were identified (Figure 4C, Table S2). The results showed that cells of clusters 2 highly expressed *Ctsd, Fxyd4*, and *Wfdc2*, which are involved in epithelial cell development and proliferation (Figure 4C-D).

**Figure 4.**
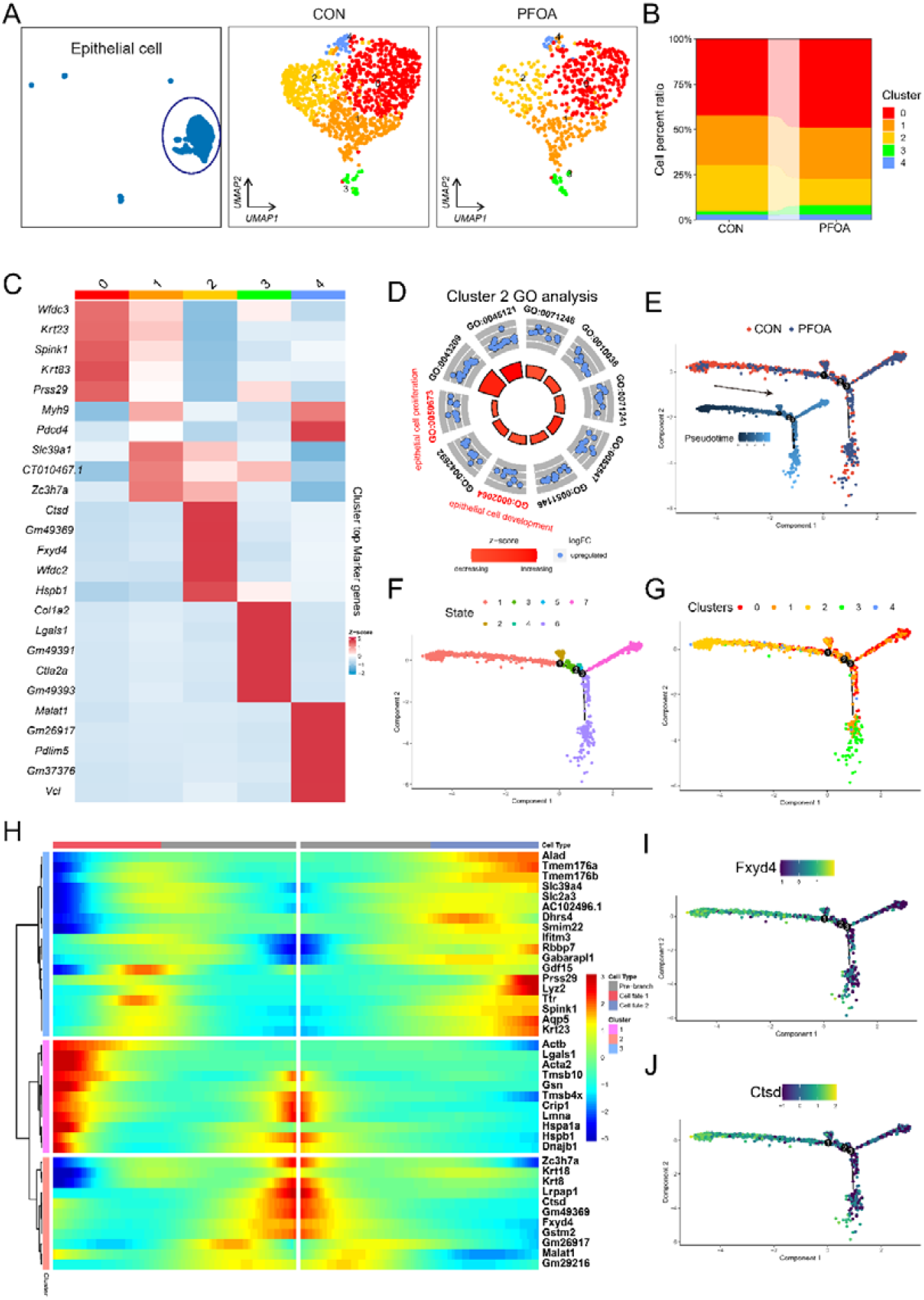
The exposure to PFOA has impaired the proportion of epithelial cell subgroups in the endometrium. (**A**) UMAP plot of 5 unbiased epithelial cell clusters of combined Control and PFOA-exposed uterus. (**B**) Percentages of the 5 epithelial cell clusters at Control and PFOA-exposed uterus. (**C**) Heatmap plot of specific marker genes of different epithelial cell clusters. (**D**) GO analysis of epithelial cell cluster 2. (**E**) Single-cell trajectories of the epithelial cell states along with the groups and pseudotime. (**F**) Single-cell trajectories of epithelial cells along with State. (**G**) Single-cell trajectories of epithelial cells along the Seurat cluster. (**H**) Heatmap of the gene expression programs of epithelial cell with different states. The gene sets were selected with qval *<* 1 × 10e-4. (**I**-**J**) Expression of *Fxyd4* and *Ctsd* along pseudotime trajectories.

To dissect the fate determination of epithelial cells, these cells were ordered along pseudotime trajectories according to gene expression (Figure 4E). Cell pseudotime trajectories revealed 7 stromal cell states and 3 branch points (Figure 4F). Clusters 2 was at the starting point of the root state (Figure 4G). As shown in Figure 4H-J, the *Fxyd4* and *Ctsd* genes were mainly expressed in state roots.

### Exposure to PFOA caused alterations in uterine metabolic status

To further analysis the changes in metabolic state, we compared the metabolites of uterus between Control and PFOA group using LCMS/MS (Figure 5A). We identified 19 up regulated metabolites and 3 down regulated metabolites (Figure 5B-C). The different metabolites were associated with many pathways like retrograde endocannabinoid signaling, linoleic acid metabolism or glycerophospholipid metabolism (Figure 5D). Metabolism analysis of scRNA data revealed different metabolites between different cell types, PFOA expose, and subclusters of epithelial cells (Figure 5E).

**Figure 5.**
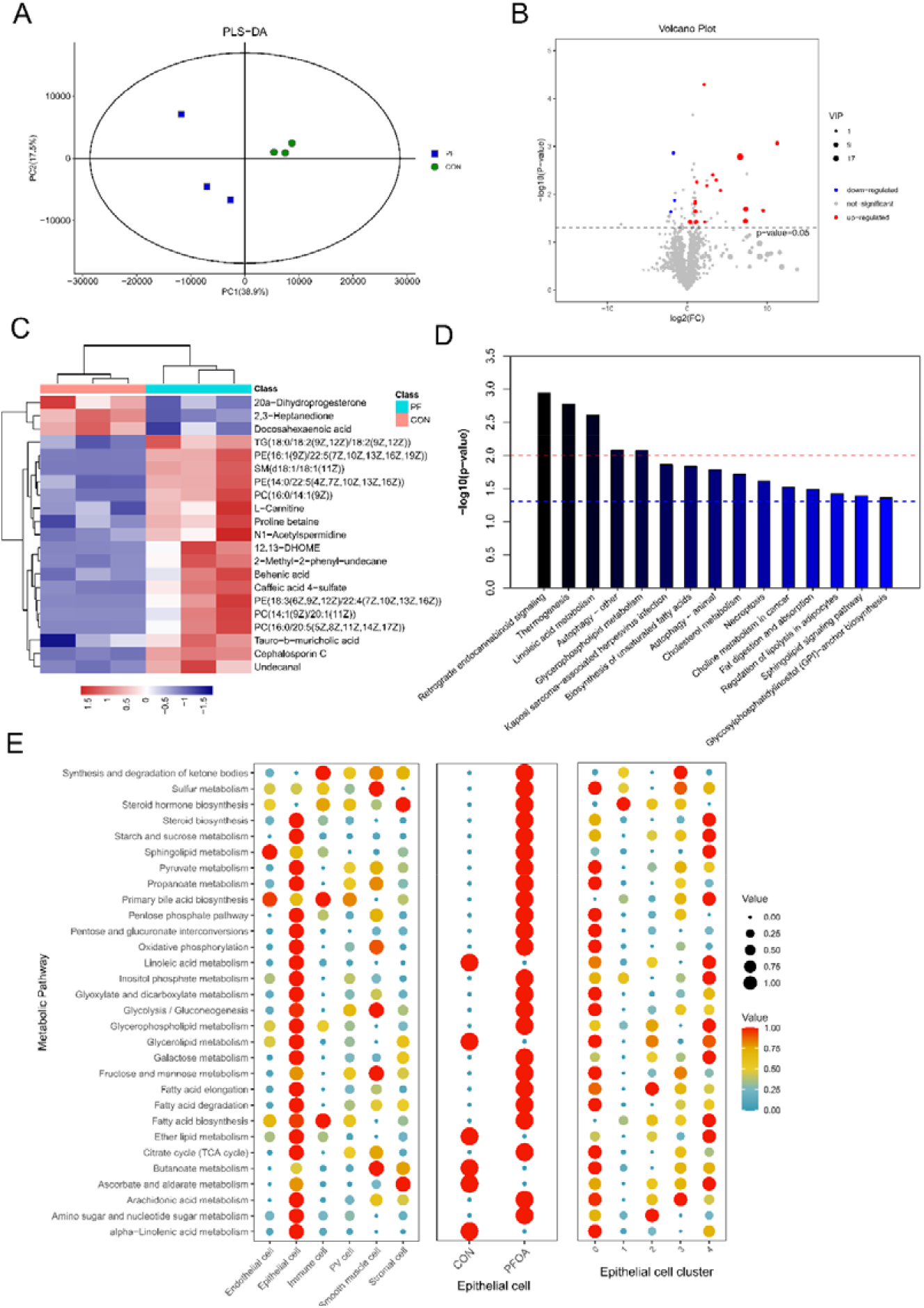
Exposure to PFOA caused alterations in uterine metabolic status. **(A)** PLS-DA analysis of Control and PFOA-exposed uterus. **(B)** Volcano plot of DEGs of Control and PFOA-exposed uterus. **(C)** Heatmap of differential metabolites of Control and PFOA-exposed uterus. **(D)** KEGG of differential metabolites of Control and PFOA-exposed uterus. **(E)** Metabolites pathway analysis between epithelial cell and epithelial cell subclusters of Control and PFOA-exposed uterus.

### PFOA affects the ANGPTL signaling communication between epithelial cells and stromal cells

To analyze the communication of these cell types in mouse uterus from Control and PFOA groups, we further performed CellChat analysis to identify the cell communication among these cell types. We found extensive and complex intercellular communication among various cell types and stromal cells sending the strongest outgoing signal to other celltypes and decreased when exposse to PFOA (Figure 6A). As showed in Figure 6B, the number of inferred interactions and interaction strength were decreased comparing to the PFOA group. The number of inferred interactions and strength of stromal cells to other celltypes were mainly decreased celltypes (Figure 6C). Thus, we investigated the relative information flow of stromal cells to other celltypes in the two groups, the result showed that the ANGPTL signaling pathway was the most impaired signaling when expose to PFOA. ANGPTL signaling was sent by stromal cells and epithelial cells was the celltype which received most of the ANGPTL signaling (Figure 6E). *Sdc4, Cdh5*, and *Cdh11* was the receptors and mainly expressed in the epithelial cells. Angptl4 was mainly expressed in stromal cells and down regulated markedly when expose to PFOA (Figure 6G-H).

**Figure 6.**
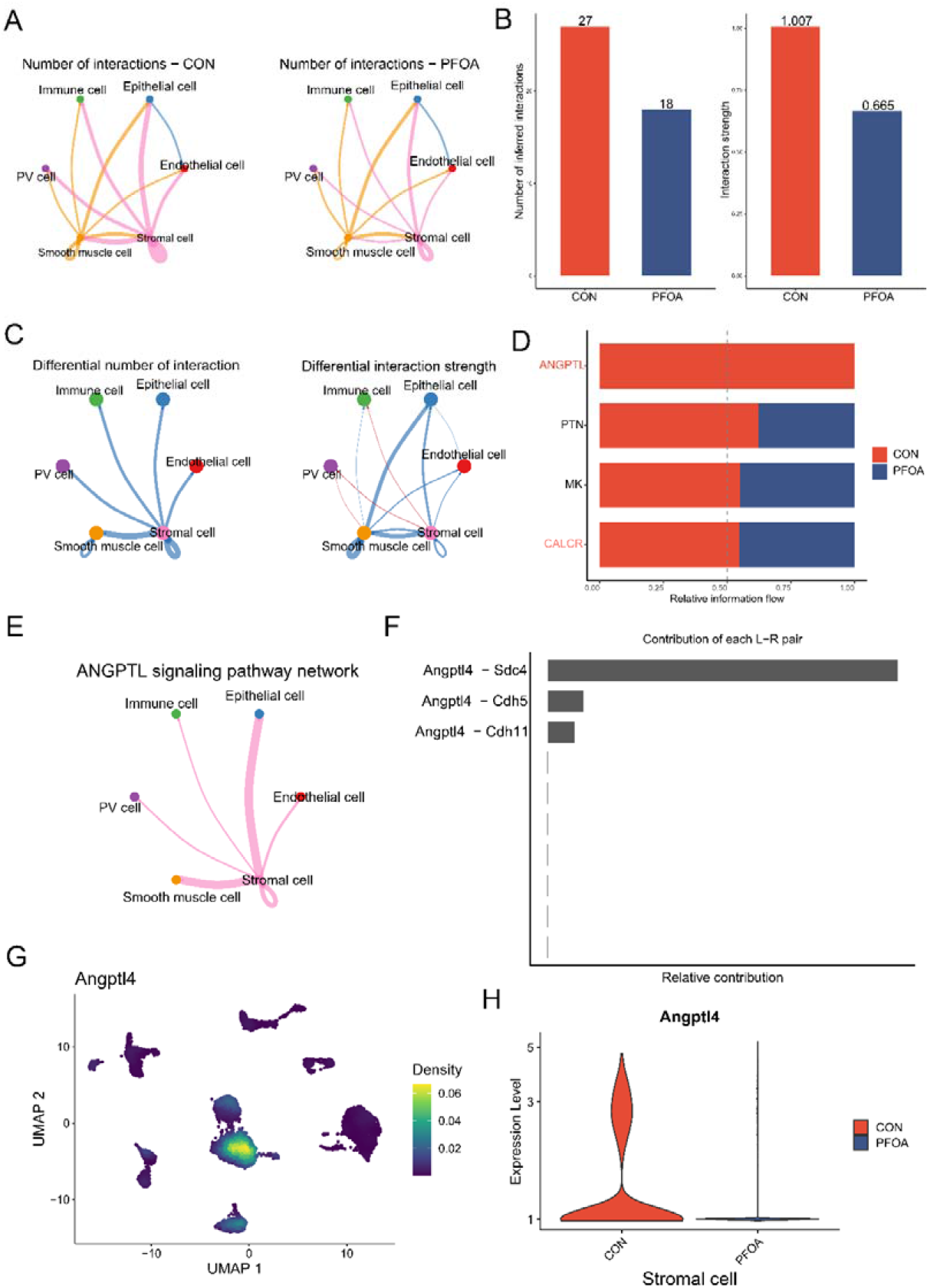
The exposure to PFOA affects the ANGPTL signaling communication between epithelial cells and stromal cells. **(A)** Cell-cell communication analysis of the six uterus cell types at Control and PFOA-exposed uterus. (**B**) Number of inferred interactions and the interaction strength at Control and PFOA-exposed uterus. (**C**) Different number of inferred interactions and the interaction strength between Control and PFOA-exposed uterus. (**D**) Relative information flow between Control and PFOA-exposed uterus. (**E**) ANGPTL signaling pathway network among the six cell types. (**F**) The contribution of each ligand–receptor (L−R) pair of the ANGPTL signaling pathway. (**G**) The expression of Angptl4 in the six cell types. (H) The expression of Angptl4 between Control and PFOA-exposed uterus stromal cells.

## Discussion

The effect of PFOA on tolerance is not much concerned by everyone. Our report expands everyone’s knowledge and helps to improve everyone’s attention. Traditional methods cannot identify the cell populations affected by PFOA. We constructed a single-cell atlas and confirmed that epithelial cells are the main affected cell type. This not only provides a meaningful dataset for related research, but also points out possible cellular targets for prevention and treatment.

The interaction between mesenchymal cells and epithelial cells is very important for embryo implantation. We found that PFOA disrupts the dialogue between them is an important mechanism.

ANGPTL4 is a secreted glycoprotein with a physiological role in lipid metabolism and a predominant expression in adipose tissue and liver. ANGPTL4 inhibits the activity of lipoprotein lipase and thereby promotes an increase in circulating triglyceride levels. Therefore, ANGPTL4 has been highly scrutinized as a potential therapeutic target. Further involvement of ANGPTL4 has been shown to occur in tumorigenesis, angiogenesis, vascular permeability and stem cell regulation, which opens new opportunities of using ANGPTL4 as potential therapeutic targets for other pathophysiological conditions^13^. ANGPTL4 was expressed in ovarian granulosa cells (GCs) and it is association with polycystic ovary syndrome (PCOS)^14^. Decreased ANGPTL4 impairs endometrial receptivity during peri-implantation period in patients with recurrent implantation failure. G protein-coupled estrogen receptor stimulates human trophoblast cell invasion via YAP-mediated ANGPTL4 expression^15^.

The impact of PFOA on the implantation rate of IVF-ET is still controversial in terms of clinical research results, which may be related to the sample size and the region of the patient^3161718^. However, the PFOA-exposed uterus model in mice revealed a role, which should arouse clinical attention to PFOA-exposed patients.

## Materials and Methods Mice

Mice were raised under specific-pathogen-free (SPF) conditions at the Experimental Animal Center of Nanjing Drum Tower Hospital (Nanjing, China). CD-1 mice were housed in a controlled environment with a 12-hour light-dark cycle, 18-20°C temperature, and 50-70% humidity. The PFOA group received PFOA (MACKLIN, Shanghai, China) at a dose of 2 mg/kg/day in the drinking water for 30 days to model PFOA exposure, while the control group received an equal volume of sterile water. The selected PFOA dose was based on previous studies. All animal experiments were conducted in accordance with the guidelines of the Laboratory Animal Management Committee (Jiangsu Province, China) and approved by the Institutional Animal Care and Use Committee of Nanjing Drum Tower Hospital (SYXK 2019-0058).

### Single-cell libraries and sequencing

The scRNA-seq library of uterus cells was constructed following the BMKMANU scRNA-seq protocol. Briefly, cell suspensions were captured on a DG1000 system (BMKMANU, Biomarker), and single-cell mRNA libraries were generated using BMKMANU DG1000 Library Construction Kits, according to the manufacturer’s protocol. The amplified cDNA libraries were then fragmented and sequenced on an Illumina NovaSeq 6000 (Illumina, San Diego). The FASTQ files were processed using BSCMATRIX to generate a single-cell feature–barcode matrixes.

### scRNA-seq data processing and analysis

The data matrixes were loaded in R (version 4.1) using the Seurat package (version 4.1.0)^19^. The Seurat object was created based on filtering parameters of “nFeature_RNA” > 200 and “nFeature_RNA” < 5000, and “percent.mt” < 25. Then, multiple samples were integrated using the “Harmony” package ^20^. After normalization and scaling, cell clustering was performed using the uniform manifold approximation and projection (UMAP) method, which is a visual method for clustering high-dimensional transcriptomic data^21^. A series of commands were executed to visualize cell clusters, including the “RunUMAP” function with a proper combination of “resolution” and “dims.use,” and “FindNeighbors” and “FindClusters” functions. To identify canonical cell cluster marker genes, the “FindAllMarkers” function and COSG package were used to identify conserved marker genes in clusters with default parameters ^22^.

### Pseudotime analysis of single-cell transcriptomes

Cell lineage trajectories were constructed using the Monocle2 package^23^. A new dataset for Monocle object was created with the gene count matrix as input, and then the “reduceDimension” and “orderCells” functions were executed to generate the cell trajectory based on pseudotime.

### Superovulation and oocyte collection

Superovulation was induced in the mice by intraperitoneal injection of 10 IU pregnant mare serum gonadotropin (PMSG) (Sansheng Pharmaceuticals, Ningbo, China) after modeling. After 48 hours, the mice were injected with 10 IU human chorionic gonadotropin (hCG) (Sansheng Pharmaceuticals, Ningbo, China). Cumulus-oocyte complexes (COCs) were collected from the ampulla of the fallopian tube 13-15 hours after the injection of hCG^24^. The MII stage oocytes were counted under a microscope after digestion with 1% hyaluronidase (Irvine Scientific, CA, USA).

### *In vitro* fertilization (IVF)

To achieve IVF, the sperm were released by cutting the epididymis tails in human tubal fluid (HTF) medium for capacitation. Cumulus-oocyte complexes (COCs) were obtained from the ampulla via superovulation and placed in 50 µL of HTF medium until insemination. After sperm capacitation, 10 µL of HTF medium containing spermatozoa was added to droplets containing COCs. The oocytes were recovered from the droplets of HTF medium 2 hours after insemination and washed 3 times in KSOM medium to remove residual GCs and spermatozoa around oocytes^25^.

The developing embryos were observed and evaluated for progression at 9 hours after insemination for zygotes, 24 hours for 2-cell embryos, 48 hours for 4-cell embryos, 72 hours for morula embryos, and 96 hours for blastocyst embryos. The formation ratios of 2-cell, 4-cell, morula, and blastocyst embryos were determined.

### Statistics of embryo implantation site

At 4.5 days post coitus (dpc), mice were sacrificed and injected with Chicago Sky Blue (Yuanye Biotechnology, Shanghai, China) intravenously. The blue staining sites were considered as the embryo implantation site^12^.

### RNA-seq and data analysis

Total RNA was extracted from ovaries using TRIzol reagent. Then, 1 µg of RNA was used to construct a cDNA library using the VAHTS Universal V6 RNA-seq Library Prep Kit for Illumina (NR604, Vazyme, Nanjing, China). The library was generated on a cBot Cluster Generation System and sequenced on the Illumina NovaSeq platform to generate 150-bp paired-end reads.

Differentially expressed genes (DEGs) were evaluated using the DEseq2^26^ packages in Bioconductor (version 1.32.0), and the DEGs were obtained using |log2-fold change|□≥□0.58 and P□≤□0.05 as cutoff criteria. Gene Ontology (GO) enrichment analysisof the large gene set (>100 genes) were performed with ClusterProfiler^27^ to detect gene-related biological processes and signaling pathways with a threshold value of “pvalueCut off = 0.05,” and top terms were displayed.

### Quantitative real-time PCR (qRT□PCR)

2 μg of total RNA was reverse transcribed into cDNA using the PrimeScript RT reagent kit (Abm, Richmond, BC, Canada). The resulting cDNA was then diluted to 200 ng/µL. Quantitative real-time PCR was performed on a qTOWER³ real-time PCR thermal cycler (Analytik, Jena, Germany) using a SYBR green PCR kit (Vazyme). The relative mRNA expression levels of *Myd88, Cldn3, Flnb Rasn* were detected and normalized to the mRNA levels of *Actb* in mice. The PCR conditions were as follows: 95 °C for 30 s (step 1), 95 °C for 10 s and 60 °C for 30 s repeating the cycle 40x (step 2). The qPCR data were analyzed using the 2-ΔΔCT method. A list of primer sequences for the genes detected is provided in Table 1.

### Spectrometry analysis of uterus

Metabolite analysis of the uterus was performed using LC/MS. Uterus samples were collected and sent to OE Biotech Co., Ltd. (Shanghai, China) for untargeted metabolome analysis. The analysis was conducted using a Nexera UPLC ultrahigh-performance liquid phase tandem QE high-resolution mass spectrometer.

### Statistical analysis

All statistics were compared using Student’s t test or one-way ANOVA with GraphPad Prism software (version 9.1), and the data are presented as the mean ± SEM from at least three replicate experiments. A p-value of less than 0.05 was considered statistically significant in this study.

## References

(1) Ding, N.; Harlow, S. D.; Randolph Jr, J. F.; Loch-Caruso, R.; Park, S. K. Perfluoroalkyl and Polyfluoroalkyl Substances (PFAS) and Their Effects on the Ovary. Hum. Reprod. Update 2020, 26 (5), 724–752. https://doi.org/10.1093/humupd/dmaa018.

(2) Rickard, B. P.; Rizvi, I.; Fenton, S. E. Per- and Poly-Fluoroalkyl Substances (PFAS) and Female Reproductive Outcomes: PFAS Elimination, Endocrine-Mediated Effects, and Disease. Toxicology 2022, 465, 153031. https://doi.org/10.1016/j.tox.2021.153031.

(3) Wang, W.; Hong, X.; Zhao, F.; Wu, J.; Wang, B. The Effects of Perfluoroalkyl and Polyfluoroalkyl Substances on Female Fertility: A Systematic Review and Meta-Analysis. Environ. Res. 2023, 216, 114718. https://doi.org/10.1016/j.envres.2022.114718.

(4) Rahman, M. F.; Peldszus, S.; Anderson, W. B. Behaviour and Fate of Perfluoroalkyl and Polyfluoroalkyl Substances (PFASs) in Drinking Water Treatment: A Review. Water Res. 2014, 50, 318–340. https://doi.org/10.1016/j.watres.2013.10.045.

(5) Zhou, Y.-T.; Li, R.; Li, S.-H.; Ma, X.; Liu, L.; Niu, D.; Duan, X. Perfluorooctanoic Acid (PFOA) Exposure Affects Early Embryonic Development and Offspring Oocyte Quality via Inducing Mitochondrial Dysfunction. Environ. Int. 2022, 167, 107413. https://doi.org/10.1016/j.envint.2022.107413.

(6) Zhang, Y.; Cao, X.; Chen, L.; Qin, Y.; Xu, Y.; Tian, Y.; Chen, L. Exposure of Female Mice to Perfluorooctanoic Acid Suppresses Hypothalamic Kisspeptin□reproductive Endocrine System through Enhanced Hepatic Fibroblast Growth Factor 21 Synthesis, Leading to Ovulation Failure and Prolonged Dioestrus. J. Neuroendocrinol. 2020, 32 (5). https://doi.org/10.1111/jne.12848.

(7) Zhang, Z.; Tian, J.; Liu, W.; Zhou, J.; Zhang, Y.; Ding, L.; Sun, H.; Yan, G.; Sheng, X. Perfluorooctanoic Acid Exposure Disrupts the Mitochondrial Electron Transport Chain in Granulosa Cells, Leading to Defect in Follicular Development. Rochester, NY July 1, 2023. https://doi.org/10.2139/ssrn.4446976.

(8) Zhou, Y.; Li, H.; Lin, C.; Mao, Y.; Rao, J.; Lou, Y.; Yang, X.; Xu, X.; Jin, F. Perfluorooctanoic Acid (PFOA) Inhibits the Gap Junction Intercellular Communication and Induces Apoptosis in Human Ovarian Granulosa Cells. Reprod. Toxicol. 2020, 98, 125–133. https://doi.org/10.1016/j.reprotox.2020.09.005.

(9) Tsang, H. Effects of Endocrine Disruptors (TCDD and PFOA) on Implantation□: An in Vitro Co-Culture Study, The University of Hong Kong, Pokfulam Road, Hong Kong SAR, 2011. https://doi.org/10.5353/th_b4717684.

(10) Mei, J.; Sheng, X.; Yan, Y.; Cai, X.; Zhang, C.; Tian, J.; Zhang, M.; Zhou, J.; Shan, H.; Huang, C. Decreased Krüppel-like Factor 4 in Adenomyosis Impairs Decidualization by Repressing Autophagy in Human Endometrial Stromal Cells. BMC Mol. Cell Biol. 2022, 23 (1), 24. https://doi.org/10.1186/s12860-022-00425-6.

(11) Zhang, Y.; Zhang, Z.; Kang, N.; Sheng, X. Generation of a Mouse Artificial Decidualization Model with Ovariectomy for Endometrial Decidualization Research. J. Vis. Exp. 2022, No. 185, e64278. https://doi.org/10.3791/64278.

(12) Tian, J.; Zhang, C.; Kang, N.; Wang, J.; Kong, N.; Zhou, J.; Wu, M.; Ding, L.; Sun, H.; Yan, G.; Sheng, X. Attenuated Monoamine Oxidase a Impairs Endometrial Receptivity in Women with Adenomyosis via Downregulation of FOXO1. Biol. Reprod. 2021, 105 (6), 1443–1457. https://doi.org/10.1093/biolre/ioab182.

(13) Li, M.; Hu, J.; Yao, L.; Gao, M. Decreased ANGPTL4 Impairs Endometrial Angiogenesis during Peri□implantation Period in Patients with Recurrent Implantation Failure. J. Cell. Mol. Med. 2020, 24 (18), 10730–10743. https://doi.org/10.1111/jcmm.15696.

(14) Jiang, Q.; Pan, Y.; Li, P.; Zheng, Y.; Bian, Y.; Wang, W.; Wu, G.; Song, T.; Shi, Y. ANGPTL4 Expression in Ovarian Granulosa Cells Is Associated With Polycystic Ovary Syndrome. Front. Endocrinol. (Lausanne). 2022, 12. https://doi.org/10.3389/fendo.2021.799833.

(15) Cheng, J.-C.; Fang, L.; Li, Y.; Thakur, A.; Hoodless, P. A.; Guo, Y.; Wang, Z.; Wu, Z.; Yan, Y.; Jia, Q.; Gao, Y.; Han, X.; Yu, Y.; Sun, Y.-P. G Protein-Coupled Estrogen Receptor Stimulates Human Trophoblast Cell Invasion via YAP-Mediated ANGPTL4 Expression. Commun. Biol. 2021, 4 (1), 1285. https://doi.org/10.1038/s42003-021-02816-5.

(16) Ma, X.; Cui, L.; Chen, L.; Zhang, J.; Zhang, X.; Kang, Q.; Jin, F.; Ye, Y. Parental Plasma Concentrations of Perfluoroalkyl Substances and In Vitro Fertilization Outcomes. Environ. Pollut. 2021, 269, 116159. https://doi.org/10.1016/j.envpol.2020.116159.

(17) Björvang, R. D.; Hallberg, I.; Pikki, A.; Berglund, L.; Pedrelli, M.; Kiviranta, H.; Rantakokko, P.; Ruokojärvi, P.; Lindh, C. H.; Olovsson, M.; Persson, S.; Holte, J.; Sjunnesson, Y.; Damdimopoulou, P. Follicular Fluid and Blood Levels of Persistent Organic Pollutants and Reproductive Outcomes among Women Undergoing Assisted Reproductive Technologies. Environ. Res. 2022, 208, 112626. https://doi.org/10.1016/j.envres.2021.112626.

(18) Hong, A.; Zhuang, L.; Cui, W.; Lu, Q.; Yang, P.; Su, S.; Wang, B.; Zhang, G.; Chen, D. Per- and Polyfluoroalkyl Substances (PFAS) Exposure in Women Seeking in Vitro Fertilization-Embryo Transfer Treatment (IVF-ET) in China: Blood-Follicular Transfer and Associations with IVF-ET Outcomes. Sci. Total Environ. 2022, 838, 156323. https://doi.org/10.1016/j.scitotenv.2022.156323.

(19) Butler, A.; Hoffman, P.; Smibert, P.; Papalexi, E.; Satija, R. Integrating Single-Cell Transcriptomic Data across Different Conditions, Technologies, and Species. Nat. Biotechnol. 2018, 36 (5), 411–420. https://doi.org/10.1038/nbt.4096.

(20) Korsunsky, I.; Millard, N.; Fan, J.; Slowikowski, K.; Zhang, F.; Wei, K.; Baglaenko, Y.; Brenner, M.; Loh, P.-R.; Raychaudhuri, S. Fast, Sensitive and Accurate Integration of Single-Cell Data with Harmony. Nat. Methods 2019, 16 (12), 1289–1296. https://doi.org/10.1038/s41592-019-0619-0.

(21) Becht, E.; McInnes, L.; Healy, J.; Dutertre, C. A.; Kwok, I. W. H.; Ng, L. G.; Ginhoux, F.; Newell, E. W. Dimensionality Reduction for Visualizing Single-Cell Data Using UMAP. Nat. Biotechnol. 2019, 37 (1), 38–47. https://doi.org/10.1038/nbt.4314.

(22) Dai, M.; Pei, X.; Wang, X.-J. Accurate and Fast Cell Marker Gene Identification with COSG. Brief. Bioinform. 2022, 23 (2), bbab579. https://doi.org/10.1093/bib/bbab579.

(23) Qiu, X.; Mao, Q.; Tang, Y.; Wang, L.; Chawla, R.; Pliner, H. A.; Trapnell, C. Reversed Graph Embedding Resolves Complex Single-Cell Trajectories. Nat. Methods 2017, 14 (10), 979–982. https://doi.org/10.1038/nmeth.4402.

(24) Sheng, X.; Yang, Y.; Zhou, J.; Yan, G.; Liu, M.; Xu, L.; Li, Z.; Jiang, R.; Diao, Z.; Zhen, X.; Ding, L.; Sun, H. Mitochondrial Transfer from Aged Adipose-Derived Stem Cells Does Not Improve the Quality of Aged Oocytes in C57BL/6 Mice. Mol. Reprod. Dev. 2019, 86 (5), 516–529. https://doi.org/10.1002/mrd.23129.

(25) Sheng, X.; Liu, C.; Yan, G.; Li, G.; Liu, J.; Yang, Y.; Li, S.; Li, Z.; Zhou, J.; Zhen, X.; Zhang, Y.; Diao, Z.; Hu, Y.; Fu, C.; Yao, B.; Li, C.; Cao, Y.; Lu, B.; Yang, Z.; Qin, Y.; Sun, H.; Ding, L. The Mitochondrial Protease LONP1 Maintains Oocyte Development and Survival by Suppressing Nuclear Translocation of AIFM1 in Mammals. eBioMedicine 2022, 75 (December 2021), 103790. https://doi.org/10.1016/j.ebiom.2021.103790.

(26) Love, M. I.; Huber, W.; Anders, S. Moderated Estimation of Fold Change and Dispersion for RNA-Seq Data with DESeq2. Genome Biol. 2014, 15 (12), 550. https://doi.org/10.1186/s13059-014-0550-8.

(27) Yu, G.; Wang, L.-G.; Han, Y.; He, Q.-Y. ClusterProfiler: An R Package for Comparing Biological Themes Among Gene Clusters. Omi. A J. Integr. Biol. 2012, 16 (5), 284–287. https://doi.org/10.1089/omi.2011.0118.

